# Dynamical consequences of context-dependent mutation: visualizing antibody sequence evolution during somatic hypermutation

**DOI:** 10.64898/2026.10.26.762018

**Authors:** Vasilisa A. Kovaleva, Hannah V. Meyer, David M. McCandlish

## Abstract

Context-dependent mutation models describe how local sequence context affects individual substitution rates, but how these biases shape sequence evolution over multiple mutational steps remains challenging to characterize. Here, we develop a framework for analyzing and visualizing long-term sequence dynamics arising from context-dependent, non-reversible mutation processes. We apply it to somatic hypermutation during antibody affinity maturation, using previously inferred context-dependent mutation rates. Our analysis recovers conventional hypermutation hot- and cold-spot patterns while exposing substantial heterogeneity in mutability within these categories. We also identify sequences with low mutability that lack conventional cold-spot motifs. Applying the framework to antigen-binding regions from different antibody-encoding gene families reveals how differences in sequence composition lead to distinct mutational trajectories through sequence space, suggesting that hyper-mutation biases constrain how these regions diversify. We further show how mutational biases defined at the level of 5-nucleotide contexts propagate to the codon and amino acid levels, producing low stationary frequencies for stop codons and tyrosine and high stationary frequencies for proline. Changes in amino acid composition between naive and memory B cells are qualitatively consistent with these predictions. Together, our results provide a general framework for connecting local context-dependent mutation rates to long-term sequence evolution and demonstrate how intrinsic somatic hypermutation biases shape the relative accessibility of different regions of sequence space during affinity maturation.

## Introduction

Single-nucleotide mutations are a key source of variation in DNA sequences. However, the rates at which these mutations occur are often context-dependent, meaning that the nucleotide mutation rates at one position depend on the nucleotide identities of possibly several flanking positions. Context-dependent mutation has been characterized through direct observations of newly arising mutations, including mutation-accumulation experiments [1] and parent–offspring trios [2], as well as through analyses of genetic polymorphism [3, 4], comparative sequence data [5], and mutational signatures in cancer [6, 7]. Previous theoretical work has also examined how context-dependent mutation shapes sequence evolution over multiple mutational steps, particularly its effects on stationary sequence composition [8]. However, understanding how these biases shape the accessibility of different regions of sequence space over time remains challenging because each mutation changes the sequence context that determines the rates of subsequent mutations. Moreover, context-dependent mutational processes need not satisfy detailed balance, a condition requiring that, at stationarity, the probability flow between any two sequences is equal in both directions.

Since detailed balance is equivalent to reversibility, such models can give rise to non-reversible Markov processes [9], which are not directly accommodated by existing approaches for visualizing evolutionary dynamics that assume reversibility [10–12]. To address these challenges, we introduce a new framework for studying sequence dynamics arising from context-dependent, non-reversible mutation processes, focusing on somatic hypermutation during antibody affinity maturation as a case study.

Antibodies are a crucial component of the adaptive immune response, important for both pathogen clearance and prevention of secondary infection. An antibody consists of two heavy chains and two light chains. Genes encoding both chains are assembled during B-cell development through a semi-random process of somatic DNA recombination, in which germline-encoded V, D (for heavy chains), and J gene segments are joined together in an imprecise manner. At the junction of recombined segments, the main antigen recognition region, complementarity-determining region 3 (CDR3), arises; the other two complementarity-determining regions, CDR1 and CDR2, are encoded in the V-gene segment. Antibodies are initially expressed on the surface of B cells as B-cell receptors (BCRs) and can later be secreted in soluble form as antibodies. Upon encountering an antigen its BCR can bind to, a B cell becomes activated and can undergo affinity maturation, a process of fine-tuning antibodies to the antigen through multiple cycles of mutation and selection. The mutational component, somatic hypermutation, is largely driven by the activation-induced cytidine deaminase, which targets sequence motifs known as hot spots, while avoiding others called cold spots [13]. Extending the hotspot–coldspot framework, numerous models have been developed to infer mutation rates and predict mutational outcomes from local sequence context, including models for all possible 5-nucleotide contexts [14–16] and, more recently, for 13-nucleotide [17] and 21-nucleotide contexts [18]. Although these contexts are much shorter than the fully reassembled BCR gene, somatic hypermutation preferentially targets relatively short regions, particularly the CDRs [16, 19] (typically up to 35 nucleotides). Such models remain essential for sequence design and function prediction: for example, they help disentangle the effects of selection and mutation during training of antibody-language models for modern antibody design [20, 21]. Here, we study the dynamical consequences of context-dependent mutation during somatic hypermutation by developing a strategy for visualizing how mutation bias shapes antibody sequence evolution over time.

Specifically, using somatic hypermutation rates previously inferred from nonproductive B-cell receptor rearrangements for all possible 5-nucleotide contexts [16], we estimate transition rates between *k*-nucleotide sequences (*k*-mers) differing by a single substitution. *k*-mer boundaries are handled by reusing flanking context from the opposite end of the *k*-mer so that every position is evaluated within a complete 5-nucleotide context. Using simulations, we confirm that for sufficiently large *k*, this approximation preserves the expected mutational dynamics. To visualize these dynamics, we extend a method originally developed for fitness landscapes [10], in which the layout reflects the evolutionary dynamics induced by the mutational process.

The resulting mutational dynamics recover the expected patterns associated with known hot- and cold-spot motifs [13]; however, they also reveal substantial variation in mutability among sequences containing the same hot- and cold-spot motifs. Furthermore, they identify sequences with low mutability that lack conventional cold-spot motifs. Extending the analysis beyond nucleotide motifs, we show that context-dependent mutation also produces systematic biases at the codon and amino acid levels, driving sequences away from stop codons and certain amino acids, such as tyrosine, while favoring others, such as proline. These biases, in turn, shape how antibodies explore sequence space during affinity maturation. We show that differences in codon composition among gene segment families result in distinct mutational trajectories for CDR1 and CDR2. Thus, these findings suggest that intrinsic somatic hypermutation biases shape not only which mutations occur, but also which regions of sequence space are more accessible than others during antibody diversification.

## Results

### Dynamics and visualization of context-dependent mutation

#### Analysis of mutational Markov chains

Consider a mutation rate matrix *M*, where *M*_*i,j*_ is the mutation rate from sequence *i* to sequence *j*. Let the total mutation rate at sequence *i*, i.e. its mutability, be 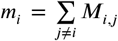. We set the diagonal entries *M*_*i,i*_ = −*m*_*i*_ so that each row of *M* sums to 0. The corresponding transition-probability matrix is then *P* (*t*) = *e*^*tM*^, where *P*_*i,j*_ (*t*) is the probability that a sequence *i* is converted by the mutational process to sequence *j* at time *t*. We assume that the mutational dynamics can in principle convert any sequence to any other sequence by some succession of mutational events. Consequently, the mutational process has a unique stationary distribution *π* such that *π*^*T*^ *P* (*t*) = *π*^*T*^ for all *t. P*_*i*_(*t*), the *i*-th row of *P* (*t*), gives the probability distribution over sequences at time *t* for a process starting from sequence *i*. Regardless of the initial sequence *i, P*_*i*_(*t*) converges to *π*^*T*^ as *t* → ∞.

The key fact that we will use to understand the dynamics of the mutational process is that for any *M* that is not a defective matrix (i.e. which has a complete set of eigenvectors), *P*_*i*_(*t*) can be expressed as a collection of deviations from the stationary distribution, each decaying at a characteristic rate. Specifically, we can write:

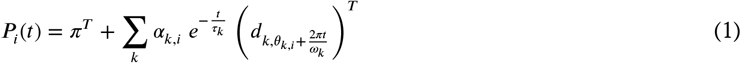

where *τ*_*k*_ is the time-scale for the decay of the *k*-th deviation, and the deviations are ordered by their decay times (*τ*_1_ ≥ *τ*_2_ ≥ *τ*_3_ …). Each deviation 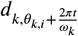 is a one-dimensional family of vectors whose entries sum to 0. This family is indexed by an angular parameter in radians, such that *d*_*k,θ*_ = *d*_*k,θ*+2*π*_ for all *θ*, and *ω*_*k*_ is the period of rotation. Finally, while the deviations themselves, their periods *ω*_*k*_, and their time-scales of decay *τ*_*k*_ are shared across starting sequences, the choice of starting sequence determines the initial weight of the *k*-th deviation *α*_*k,i*_, as well as an initial phase for the *k*-th deviation *θ*_*k,i*_.

The way to understand Equation 1 is to consider how *P*_*i*_(*t*), initially concentrated entirely at sequence *i*, diffuses across sequence space and eventually approaches the stationary distribution *π*. The strongest barriers to diffusion have the largest *τ*_*k*_ and create long-term excesses and deficits of probability in the regions of sequence space described by the most slowly decaying deviations from stationarity. Because *α*_*k,i*_ for small *k* describe the contributions of these slowly decaying deviations, using the *α*_*k,i*_ and the corresponding initial phases *θ*_*k,i*_ as embedding coordinates places together sequences for which *P*_*i*_(*t*) approaches *π*^*T*^ in a similar way. Intuitively, the resulting clusters represent sets of sequences separated by barriers that are the most difficult for the diffusion process to cross.

At a more technical level, the elements of Equation 1 can be derived from the eigendecomposition of *M*. Specifically, the eigenvalues of *M* are either real or complex. For left and right eigenvectors *l* and *r* of *M* with eigenvalue *λ*, we normalize *r* such that *r*^∗^*D*_*π*_*r* = 1 and *l* such that *l*^∗^*r* = 1, where *D*_*π*_ is the diagonal matrix with diagonal entries given by the vector of stationary probabilities and the star indicates the conjugate transpose. For matrices *M* with repeated eigenvalues, we use the fact that *M* is non-defective to choose bi-orthogonal left and right eigenbases, such that each left eigenvector is orthogonal to every right eigenvector except for the one it corresponds to.

The dominant eigenvalue equals 0, with left eigenvector *π*^*T*^ and right eigenvector consisting of all 1’s (because all rows of *M* sum to 0). Complex eigenvalues come in conjugate pairs: if *λ* = *a* + *bi*, then the paired eigenvalue is *a* − *bi*, and the entries of the corresponding eigenvectors are also complex conjugates. Each non-zero real eigenvalue and each complex-conjugate pair contributes one term to the sum in Equation 1. The corresponding decay time-scale *τ*_*k*_ is given by the negative reciprocal of the real part of *λ*, i.e. 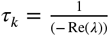. For real eigenvalues, the corresponding *d*_*k,θ*_ is independent of *θ*, such that *d*_*k,θ*_ = *l*; since in this case the phase and period are irrelevant, we simply suppress them from our notation. If *λ* is real, the corresponding *α*_*k,i*_ is simply *r*_*i*_, the *i*-th entry of the corresponding right eigenvector *r*.

For a complex-conjugate eigenvalue pair, we must define an entire one-dimensional family of deviations rather than a single deviation. By convention, we use the eigenvectors *l* and *r* associated with the member of the pair having a positive imaginary part, *λ* = *a* + |*b*| *i*. We then define *d*_*k,θ*_ = 2 (Re(*l*) cos *θ* + Im(*l*) sin *θ*), which gives the real-valued deviation associated with the complex eigenvector *l* at phase *θ*. For a process starting from sequence *i*, the initial weight and phase are given by the magnitude of *r*_*i*_ (*α*_*k,i*_ = |*r*_*i*_|), and the angle between *r*_*i*_ and the positive real axis of the complex plane (*θ*_*k,i*_ = Arg(*r*_*i*_)), respectively.

#### Construction of visualizations

To construct a visualization based on Equation 1, we want to represent the weights of the most slowly decaying deviations from stationarity, i.e., those associated with the longest timescales, *τ*_1_, *τ*_2_, …. We thus construct a set of embedding coordinates, which, following Martí-Gómez *et al*. [12], we call diffusion axes, starting with *τ*_1_, and then moving sequentially through the *τ*_*k*_. Each deviation contributes one dimension to the visualization if the corresponding eigenvalue is real, and two dimensions if it is complex. To encode the time-scale of each deviation, we multiply the corresponding weights by 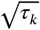, so that squared Euclidean distances in the resulting visualizations have units of time. Thus, for a real eigenvalue *λ* with eigenvectors *l* and *r* associated with *τ*_*k*_, the corresponding diffusion axis is 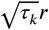, so that each initial weight is multiplied by 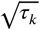. For a complex-conjugate eigenvalue pair, we want to represent each sequence at 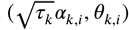 in polar coordinates, corresponding to 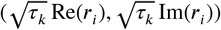 in Cartesian coordinates. For simplicity, we label the real and imaginary components associated with a complex-conjugate eigenvalue pair as the real and imaginary diffusion axes *k*, respectively, so that the axis number always corresponds to the time-scale *τ*_*k*_, regardless of how many preceding eigenvalues were real or complex. In the special case where *M* is the rate matrix of a reversible chain, all eigenvalues are real, and the right eigenvectors can be chosen to be *D*_*π*_-orthogonal. Then, the method reduces to that of [10] as implemented in [12]. Our treatment extends this visualization method to any non-defective rate matrix *M*, and may also be useful in the defective case, provided that the missing eigenvector is not associated with the few slowest time-scales used in the visualization.

Although in the reversible case the visualizations have many provable mathematical properties [10, 12], these are for the most part lost in the non-reversible case. Nonetheless, a strong relationship between the visualizations and the evolutionary dynamics remains: the expected position of the population in the visualization, averaged over its possible states according to the probability distribution *P*_*i*_(*t*), moves in a highly regular way as *t* increases. In particular, along a diffusion axis corresponding to a real eigenvalue, the expected position of the population approaches the origin exponentially at rate 1/*τ*_*k*_. For the real and imaginary embedding axes corresponding to a complex-conjugate eigenvalue pair, the expected position spirals towards the origin, with exponential decay rate 1/*τ*_*k*_ and period *ω*_*k*_.

Importantly, for our application to affinity maturation, it is quite clear that the mutational process remains far from its stationary distribution during natural B-cell maturation. In our model, the slowest deviation from stationarity decays on a characteristic timescale of *τ* = 1.77, corresponding to approximately 15 mutations per 9-mer. When normalized by sequence length, this far exceeds the number of mutations typically accumulated during affinity maturation (20-24 across an entire heavy chain variable region [22], which is approximately 300-400 nucleotides long). Nevertheless, the stationary frequencies are useful to examine because they characterize the long-term tendencies of the mutation process; we often include them as an additional plotting coordinate to highlight which sequences are favoured or depleted by the mutational process. The diffusion axes complement this information by characterizing the barriers to movement through sequence space, thereby revealing which regions of sequence space are more readily accessible on the shorter timescales relevant to affinity maturation.

#### Treatment of context-dependent mutation

To construct the matrix *M* of mutation rates between *k*-nucleotide sequences, we used a model of somatic hypermutation [16] trained on reconstructed phylogenies of BCR rearrangements, from which both positional and sequence-context preferences of hypermutation targeting were inferred. To minimize the effect of selection during affinity maturation, the authors trained the model on nonproductive, out-of-frame BCR rearrangements, which do not encode functional receptors. The model did not impose reverse-complement symmetry, so that the inferred mutation rates can differ between the corresponding mutations in reverse-complementary sequence contexts. The resulting context model estimated mutation rates for all possible 5-nucleotide motifs, with the target nucleotide located in the middle and therefore in the context of the 2 preceding and 2 subsequent nucleotides.

Importantly, to construct *M*, we need to calculate mutation rates at every position of each *k*-mer, including positions for which the full sequence context extends beyond the boundaries of the *k*-mer. To address this problem, we applied a “padding” procedure that is equivalent to imposing a periodic boundary condition: when the target nucleotide is located near one end of the sequence, the missing flanking context is taken from the opposite end. This allows every position to be evaluated within a complete 5-nucleotide context (see Methods for additional details). Since we are primarily interested in coding sequence evolution, for codon-based analyses we defined the middle 3 nucleotides of the *k*-mer to be the codon, where we define the right-hand nucleotide of the two central positions as the codon center when *k* is even. For stationary frequencies, codon-level values were calculated by summing the stationary frequencies of all *k*-mers containing that codon at the central position, and for diffusion axes, codon-level values were calculated by averaging the corresponding eigenvector components across all *k*-mers sharing the same central codon.

Clearly, for small *k*, this padding procedure can introduce artifactual dependencies, but such artifacts will tend to decrease as the sequence length increases. On the other hand, increasing sequence length substantially expands the state space and computational complexity. To determine the optimal sequence length, we therefore constructed mutation rate matrices for *k*-mers with *k* = 5 … 10 and compared their stationary distributions and first diffusion axes across successive values of *k*. Since the dimensionality of the eigenspace depends on the number of possible *k*-mers, direct comparison between different *k* is not possible, so instead for each *k* we used values for the central codon of each *k*-mer.

As expected, padding of short sequences (*k* = 5) produced spurious eigenvector values arising from context-overlap artifacts (Extended Data Fig. 1a-b, first column). However, as *k* increased, eigenvectors derived from transition matrices with successive *k* became progressively more consistent, indicating approximate convergence of the resulting mutational dynamics (Extended Data Fig. 1a-b, second and third columns). To further validate the stationary frequencies inferred from the transition matrices, we performed Gillespie simulations on 2000 random BCR sequences of naive B cells from the Observed Antibody Space database [23]. After *t* = 300 (much longer than the characteristic time scale of relaxation to stationarity, *τ*_1_ = 1.77), we measured the resulting 5-mer frequencies and compared them with the stationary 5-mer frequencies predicted by the *k*-mer model. The agreement between simulated and theoretical frequency distributions increased with greater *k* (Extended Data Fig. 1c). Based on these analyses, we decided to proceed with *k* = 9 as an appropriate compromise between accuracy and computational tractability.

### Somatic hypermutation biases shape the accessibility structure of sequence space

Somatic hypermutation biases are traditionally described in terms of hot- and cold-spot motifs [13]. By visualizing the mutational dynamics of all possible 9-mers, we were able to recover the expected pattern (Figure 1a) for sequences containing these motifs. On average, 9-mers containing a hot-spot motif (WRC) exhibited lower stationary frequencies, indicating that these sequences are preferentially depleted by the mutation process. The 9-mer with the lowest stationary frequency contained multiple hot-spot motifs (WRC [24] and WA [25], Figure 1a, left panel inset). Similarly, 9-mers containing a cold-spot motif (SYC [26]) exhibited higher stationary frequencies. However, the stationary-frequency distributions of hot- and cold-spot-containing sequences overlapped substantially, highlighting the limited specificity of these motif definitions. We also identified sequences with low mutability that lack conventional cold-spot motifs, including poly-G and poly-T sequences (Figure 1a, right). Of note, although substitution rate and calculated stationary frequencies were clearly related, the relationship was not exact (Extended Data Fig. 2a), indicating that stationary frequencies capture properties of the mutational dynamics beyond just the mutability of individual sequences. Also, as expected from the strand asymmetry of somatic hypermutation [27], complementary sequences exhibited different stationary frequencies (Extended Data Fig. 2b).

**Figure 1.**
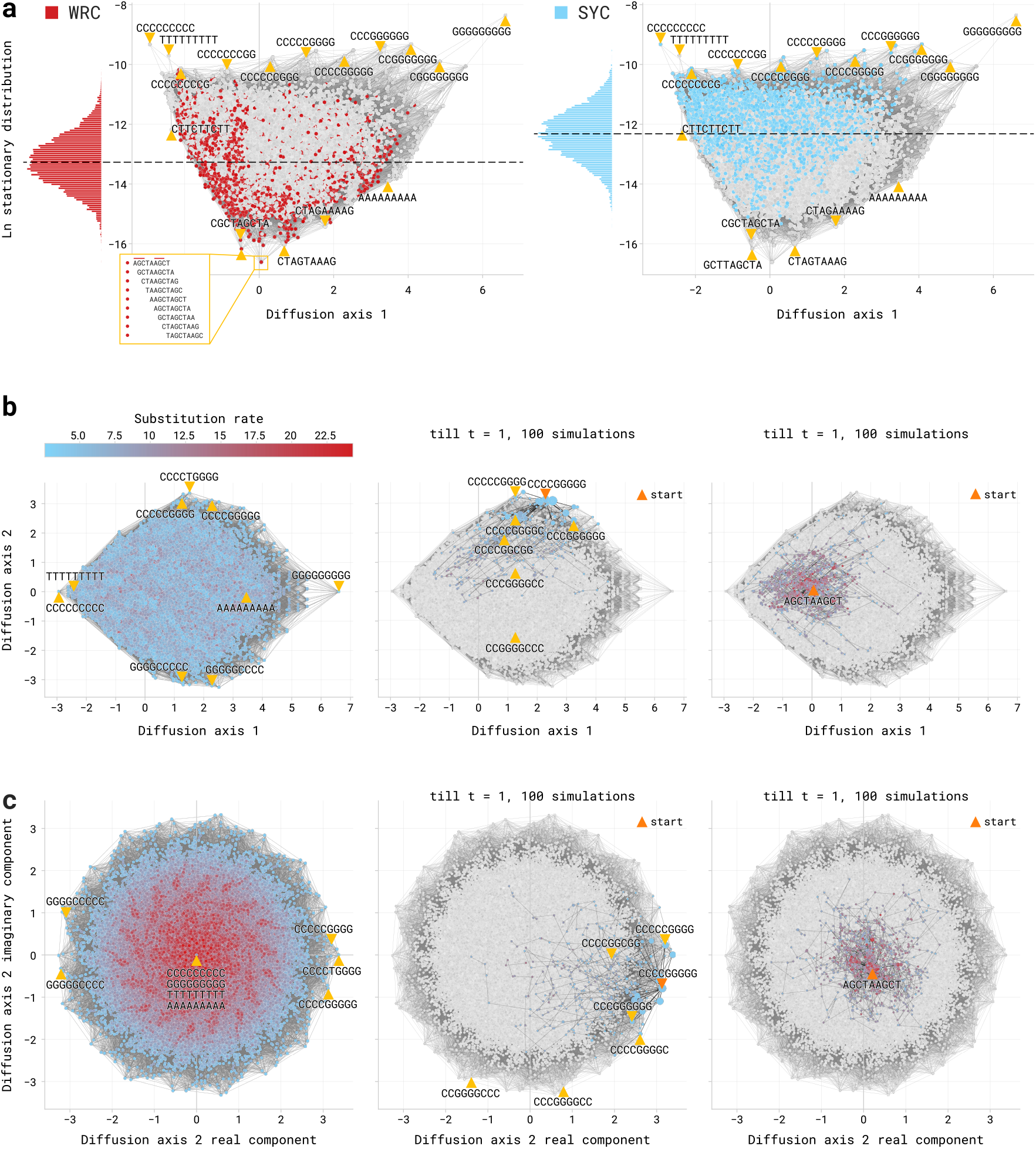
Mutational dynamics of 9-mers. **(a)** Distribution of all possible 9-mers along diffusion axis 1 and their natural log stationary frequencies. Sequences containing hot-spot (WRC, red, left) or cold-spot (SYC, blue, right) motifs are highlighted. Marginal histograms show the distribution of log stationary frequencies for motif-containing sequences. Dashed horizontal lines show the median of the corresponding distribution. The inset demonstrates how each plotted point represents cyclic rotations of the same sequence, which all share the same stationary frequency and the same position along diffusion axis 1. **(b)** Distribution of all possible 9-mers along diffusion axis 1 and the real component of diffusion axis 2. Left: 9-mers are colored by their total substitution rate. Middle and right: 100 simulated mutation trajectories, each run until *t* = 1 from starting sequences CCCCGGGGG (middle) and AGCTAAGCT (right). **(c)** Same as in (b), but using real and imaginary components of the diffusion axis 2. **(a-c).** Edges connect sequences differing by a single-nucleotide substitution.

While stationary frequencies describe the system’s equilibrium, diffusion axes (derived from right eigenvectors) characterize how sequences approach that equilibrium, with diffusion axis 1 reflecting the most slowly decaying deviation from stationarity. Consequently, the sequences separated along diffusion axis 1 are relatively inaccessible from one another via somatic hypermutation. For example, poly-T and poly-C sequences are located near each other, but poly-G and poly-C are strongly separated (Figure 1a-b), indicating that the mutational accessibility cannot be approximated simply by the number of substitutions.

To visualize mutational walks in sequence space, we employed Gillespie simulations (Figure 1b-c, second and third column). We observed that when the process starts from a 9-mer exhibiting a low substitution rate (CCCCGGGGG), visited sequences accumulate in a small number of nodes (Figure 1b-c, second column), while starting from a hot-spot-motif-enriched sequence (AGCTAAGCT, Figure 1b-c, third column) results in a more diffuse distribution of the visited sequences. Nonetheless, both distributions remain relatively localized in the visualizations, consistent with the interpretation that sequences widely separated in the visualizations are relatively difficult to reach from one another through mutation.

Mutation matrices describing context-dependent processes can violate detailed balance (i.e. can be non-reversible), allowing for complex-conjugate eigenvalues and eigenvectors, as observed in our model (Figure 1c). Intuitively, a lack of reversibility implies the presence of cyclic probability flow between sequences. The imaginary parts of the eigenvalues correspond to oscillatory components of the dynamics and therefore describe the timescale of rotation of the expected position of the population in the visualization. In our example, however, the calculated periods of rotation for the most slowly decaying deviations from equilibrium are all roughly 2-4 orders of magnitude larger than the characteristic time scale of decay for the corresponding deviation from stationarity (Extended Data Fig. 2c), meaning that the rotation occurs much more slowly than relaxation toward the stationary distribution, and therefore can be neglected in practice over the time-scale of this relaxation.

Visualization using real and imaginary components of the second diffusion axis (Figure 1c) shows that the sequences that are far from the origin, and hence diffuse in a manner that is strongly influenced by this deviation from stationarity, often contain combinations of poly-G and poly-C runs (Extended Data Fig. 2d). Intuitively, these runs are very cold, and so they are hard to remove, making it relatively difficult for mutations to transform one arrangement of poly-G and poly-C runs into another, such as a cyclically shifted arrangement of the same sequence. Interestingly, although these sequences with poly-G and poly-C runs generally have low substitution rates, they are not among the “coldest” sequences. This observation can be explained by the relatively high mutability of the G-C boundary: although poly-C or poly-G runs are indeed “cold”, the boundary between G and C stretches is hotter, resulting in somewhat lower stationary frequencies of intermediate sequences containing both G and C (Figure 1a) as compared to the fully homomeric poly-C or poly-G sequences.

### Somatic hypermutation shapes codon and amino acid composition

The differences in stationary frequencies among 9-mers were substantial: the most frequent 9-mer (poly-G sequence) was estimated to occur more than 3,000 times as often as the least frequent one (AGCTAAGCT, which consists of multiple hot-spot motifs). In contrast, when we summed all the stationary frequencies for the central codon in the 9-mer, these differences became considerably smaller (Figure 2a, a 7-fold difference between the most frequent GGG and least frequent TAA codons). Similarly, differences in stationary frequencies were again within a roughly 10-fold range at the amino acid level (Figure 2b).

**Figure 2.**
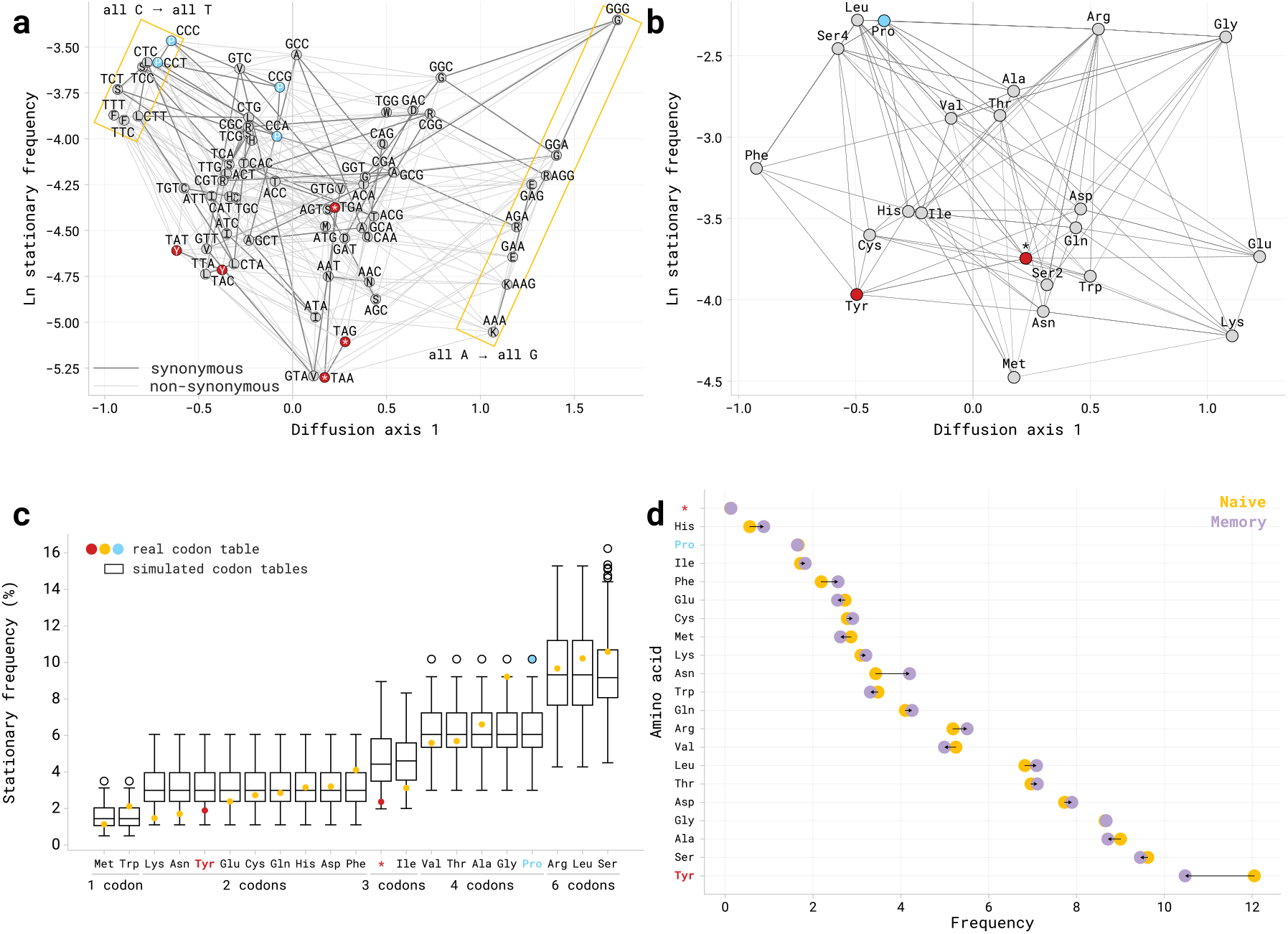
Context-dependent mutation biases propagate to codon and amino-acid levels. **(a)** Visualization of all possible codons along diffusion axis 1 and their log stationary frequencies. Edges connect codons differing by a single-nucleotide substitution; dark edges mark synonymous substitutions, and light edges mark non-synonymous substitutions. To obtain diffusion-axis values for central codons, we averaged the diffusion-axis values for multiple 9-mers with a given central codon. **(b)** The same as in (a), but at the amino acid level. Amino acids that are mutually accessible via a single nucleotide substitution are connected by edges; the two disconnected serine codon blocks (Ser2 and Ser4) are treated as if they coded for separate amino acids. **(c)** Stationary frequencies of amino acids under the standard genetic code (colored points) compared with distributions obtained from simulated codon tables (boxplots). Amino acids are sorted by codon degeneracy. **(d)** Amino acid frequencies in CDR3 sequences from naive (yellow) and memory (purple) B cells. **(a-d)** Tyrosine (red), proline (blue), and stop codons (red stars) are highlighted.

We observed no clear pattern separating synonymous from non-synonymous substitutions between codons (Figure 2a). Instead, the codons were spread roughly according to their purine vs pyrimidine composition along diffusion axis 1, and their stationary frequencies reflected their C/T and G/A ratios. Among all codons, stop codons had particularly low stationary frequencies, especially TAA and TAG. Similarly, tyrosine codons had relatively low stationary frequencies. This suggests that somatic hypermutation is biased against stop codons and tyrosines, driving sequences away from them. In contrast, proline codons exhibited high stationary frequencies, suggesting that somatic hypermutation is biased toward retaining proline-encoding codons.

To assess how likely such a distribution of stationary frequencies among codons is, we performed codon-table simulations, preserving codon table degeneracy and block organization of synonymous codons (Figure 2c). The standard genetic code yielded unusually high stationary frequencies for proline codons and unusually low frequencies for stop codons and tyrosine codons, with only 6.25%, 1.22%, and 15.62% of simulated codes, respectively, producing equally or more extreme frequencies.

To examine whether the predicted trends are reflected in observed antibody repertoires, we compared the amino acid frequencies in CDR3, which is targeted the most during somatic hypermutation [16, 19], in BCRs from naive (before somatic hypermutation; 7,228,250 CDR3 sequences) and memory B cells (after somatic hypermutation; 8,521,052 CDR3 sequences) from 3 individuals [28] (Figure 2d). As expected, stop-codon frequency did not change and remained very low, likely because BCRs that acquire premature stop codons are eliminated during the selection step of affinity maturation. In CDR3s from naive B cells, tyrosine was the most abundant amino acid and occurred at a disproportionately high frequency of 12%, but in CDR3s from memory B cells, tyrosine frequency dropped substantially, consistent with the predicted effect of somatic hypermutation biases. In contrast, proline frequency did not change. Thus, although selection also shapes antibody sequences during affinity maturation, the observed changes in amino acid composition are qualitatively consistent with the effects of the underlying mutational biases predicted by our model.

### Somatic hypermutation drives CDR-specific exploration of sequence space across V-gene families

By visualizing mutational dynamics for all possible 9-mers, we uncovered general trends in the somatic hypermutation process. In practice, however, many 9-mers are rarely observed in BCR gene sequences (for example, a poly-T 9-mer was found in 1 sequence among 43,948 sequences from OAS [23]; see *Methods, Data*). To better understand how BCRs explore sequence space, we examined where 9-mers derived from heavy-chain V gene segments in the ImMunoGeneTics database [29] are located relative to the structures in sequence space that our analysis has uncovered so far.

Based on sequence similarity, V gene segments are grouped into gene families, where all gene segments within a family share at least 75% nucleotide identity (IMGT classification; [30]). Therefore, differences in sequence composition between families, such as codon usage (Extended Data Fig. 3), may cause somatic hypermutation biases to produce distinct long-term trajectories through sequence space. Here, we present our analysis for the IGHV1 and IGHV3 gene families, the two families with the largest numbers of gene segments.

We focused on the 9-mer composition of CDR1 and CDR2, as these regions are encoded within germline segments and, together with CDR3, are frequently mutated during somatic hypermutation [16, 19]. The 9-mers derived from CDR1 and CDR2 had broadly overlapping distributions along diffusion axis 1 in the IGHV1 family, but qualitatively distinct distributions in the IGHV3 family (Figure 3, first column). To investigate the mutational consequences of these differences, we again employed Gillespie simulations. 9-mers derived from CDR1 followed broadly similar mutational trajectories across V-gene families (Figure 3, second column). In contrast, the regions of sequence space explored by CDR2 9-mers differ substantially between the V-gene families (Figure 3, third column). Overall, these results suggest that somatic hypermutation can drive CDR2 sequences from different V-gene families to explore distinct regions of sequence space, even when CDR1 sequences follow relatively similar mutational trajectories between families.

**Figure 3.**
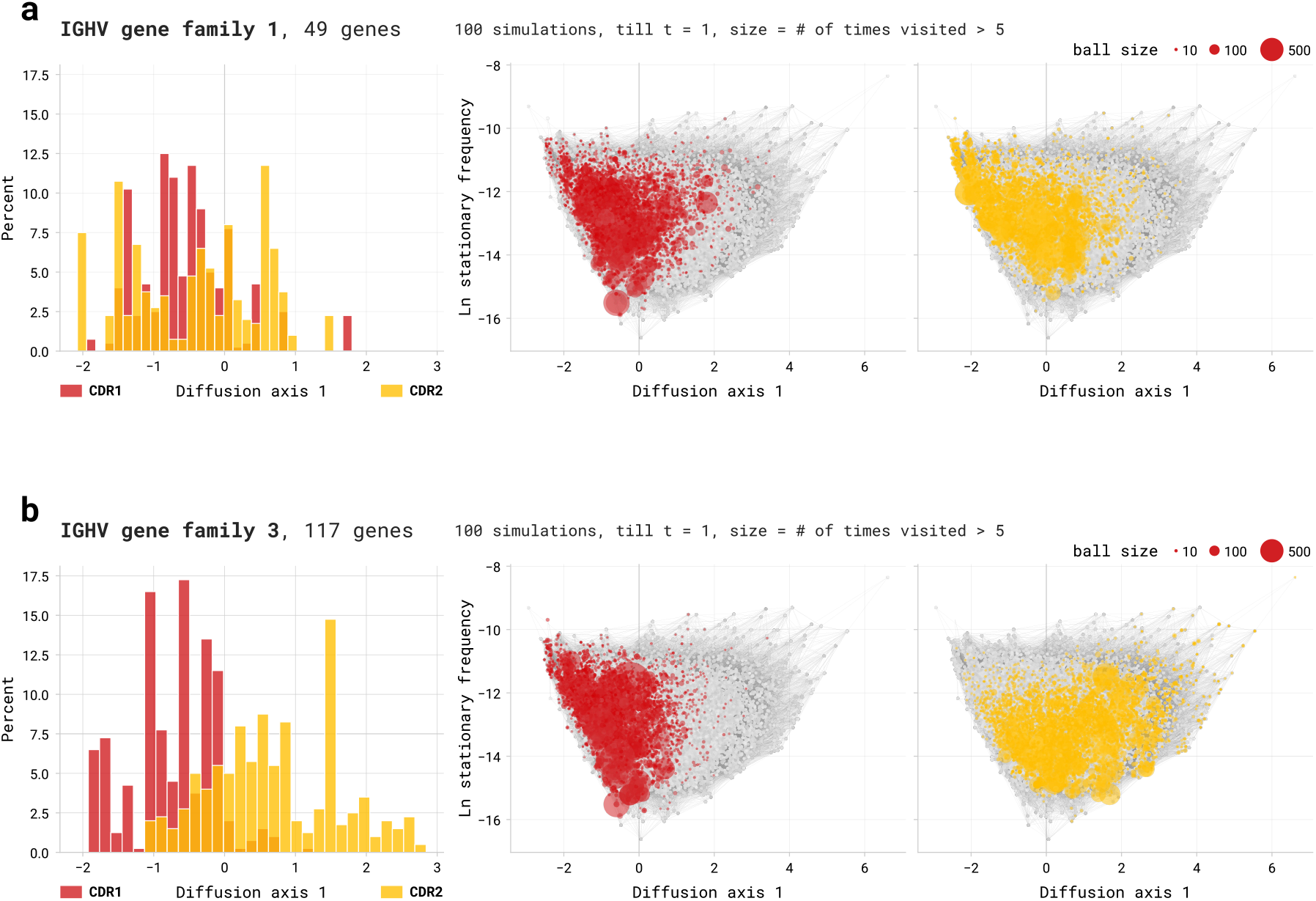
Differences in V-gene sequence composition lead to distinct mutational trajectories through sequence space. **(a)** Distribution and simulated mutation trajectories of 9-mers derived from CDR1 (red) and CDR2 (yellow) regions of V genes from the IGHV1 gene family. Left: distribution of initial 9-mers along diffusion axis 1. Middle and right: 9-mers visited during 100 simulations till *t* = 1, starting from 100 randomly sampled 9-mers derived from CDR1 and CDR2 sequences, respectively. Circle size indicates the number of visits to each 9-mer, with only sequences visited more than 5 times highlighted. **(b)** The same as in (a) for V-genes from the IGHV3 gene family.

## Discussion

Context-dependent mutation models provide estimates of mutation rates at individual sites, but understanding how these biases shape sequence evolution during multiple mutational steps remains challenging. Here, we developed a framework that enables the study of long-term mutational dynamics. To construct a mutation rate matrix for all possible sequences of an arbitrary length using a given context model, we used a “padding” procedure that provides the missing flanking context at sequence boundaries, allowing mutational rates to be estimated for every nucleotide position in the sequence. We then represented the resulting mutational dynamics as a continuous-time Markov process and characterized both the stationary distribution of sequences and their trajectories through sequence space. Importantly, our approach extends existing methods for visualizing evolutionary dynamics to non-reversible mutation processes. Although we apply the developed framework to somatic hypermutation of antibodies during affinity maturation, it can be applied to other context-dependent mutation processes, such as those occurring in the germline [2, 3], or somatic mutation during cancer progression [6, 7].

The locations of somatic hypermutation hotspots strongly constrain which mutations are accessible during affinity maturation [31], as demonstrated by beneficial but very rare mutations required for broadly neutralizing anti-HIV antibodies [32]. We used stationary frequencies and the slowest-decaying deviations from stationarity (diffusion axes) to characterize long-term mutational tendencies, revealing properties of the mutation process that cannot be explained by substitution rates alone. Using a mutation rate matrix for all possible 9-mers, our framework recapitulates conventional hot- and cold-spot patterns [24–26]. Consistent with previous findings, we show that short hot- and cold-spot motifs are insufficient to fully characterize mutational dynamics [13]. Moreover, we show that sequences with similar mutability can follow substantially different mutational trajectories. In particular, poly-C sequences contain the conventional cold-spot motif SYC, whereas poly-G sequences do not, yet exhibit similarly low mutability. These sequences are located at opposite ends of diffusion axis 1, despite differing by the same number of substitutions as poly-T and poly-C sequences, which are located close together. These results highlight poly-G sequences as an additional class of cold spots not captured by the conventional SYC motif, potentially arising from the presence of poly-C on the complementary strand but exhibiting distinct mutational trajectories, possibly reflecting strand asymmetry of somatic hypermutation targeting and repair [27, 33].

The consequences of somatic hypermutation depend not only on the mutation-targeting machinery but also on the organization of the genetic code. Thus, context-dependent somatic hypermutation biases translate into distinct mutational patterns at the codon and amino acid levels. Our framework shows that somatic hypermutation favors the enrichment of certain codons and the depletion at stationarity of others. In particular, stop codons have low stationary frequencies, which could reflect either a low probability of generating them through mutation or a high probability of mutating away from them. The former could provide an evolutionary advantage by reducing the occurrence of non-functional receptors. However, the protective effect of the latter is less apparent. Given the estimated somatic hypermutation rate of approximately 10^−3^ mutations per base pair per cell division [34], a second mutation correcting a premature stop codon would be unlikely to occur within the same division. A clearer example is the bias against tyrosine. Tyrosine is enriched in antigen-contacting regions of naive BCRs, where its aromatic side chain allows various interactions with antigens [35]. The previously described decrease in tyrosine frequency during affinity maturation [36] could contribute to the diversification and refinement of BCR binding surfaces [37]. In contrast, proline codons have high stationary frequencies, suggesting that somatic hypermutation tends to enrich proline-encoding codons. Given the strong conformational constraints imposed by proline [38], this bias might help preserve structural features during affinity maturation. Our codon-table simulations further show that the standard genetic code is among the few that produce such high stationary frequencies for proline codons. Interestingly, somatic hypermutation has also been shown to antagonize codon optimality, potentially reducing antibody expression [39]. Together, these observations suggest that antibody diversification involves trade-offs between mutational accessibility and the structural and functional properties of the resulting proteins [40].

Codon usage of antibody V genes has evolved to direct somatic hypermutation toward complementarity-determining regions [19]. More recently, DNA flexibility has been suggested as another sequence-encoded property that influences preferential somatic hypermutation targeting [41] in V genes. This raises the question of whether differences in sequence composition among V-gene families lead to distinct mutational trajectories. Indeed, we show that 9-mers derived from the IGHV1 and IGHV3 families explore different regions of sequence space during somatic hypermutation. Consistent with this finding, differences in diversification patterns between individual V-genes have previously been observed in antibody repertoires [42]. Interestingly, in our analysis, differences between gene families are particularly pronounced for CDR2, whereas CDR1 sequences follow relatively similar trajectories.

Our framework has several limitations. Most importantly, we consider only the mutation process, without accounting for selection and competition between B cell clones during affinity maturation. Nonetheless, understanding the mutational process in isolation is an important step towards understanding B cell maturation more generally. Also, while not appropriate for the specific process of B cell maturation, we do note that for applications where a weak mutation assumption is appropriate, natural selection can easily be added to our framework by incorporating the probability of fixation [10, 12]. A second set of limitations arises from the underlying mutation model. Our current implementation relies on a 5-nucleotide context model, which does not capture longer-range interactions or positional preferences in somatic hypermutation targeting [16, 43]. Nevertheless, the framework is not restricted to this context length and could be applied to more complex mutation models. The padding procedure also introduces artificial interactions between sequence boundaries, although we show that their effects decrease with increasing sequence length. Furthermore, our predictions depend on mutation rates inferred from nonproductive BCR rearrangements. Recent work has shown that models inferred from nonproductive sequences and synonymous substitutions can produce substantially different predictions [17]. Beyond the sequence-context model, we make several simplifying assumptions about the mutation process itself. We assume constant mutation rates, although somatic hypermutation activity can vary during affinity maturation [34, 44]. Recent theoretical work has shown that fitness-dependent mutation rates can substantially affect evolutionary outcomes, potentially accelerating adaptation [45]. However, changes affecting all mutation rates equally would only rescale time, without changing the structure of the mutational landscape. We also restrict individual mutation events to single-nucleotide substitutions, excluding relatively rare insertions and deletions [46]. Finally, our mathematical framework assumes that the mutation rate matrix is nondefective. However, the visualization may remain applicable to defective matrices, provided that the slowest modes used for the embedding are unaffected.

Despite these limitations, we demonstrate how the proposed framework can be used to address biological questions about sequence evolution, using somatic hypermutation as a case study. It recovers previously described mutation patterns, suggesting that the framework captures biologically relevant properties of the mutation process. Future extensions could incorporate more complex context-dependent mutation models and account for selection. Importantly, the framework is not restricted to antibody evolution and could be applied to other biological processes with pronounced context-dependent mutation biases. More generally, our approach provides a way to move from local mutation rates to the long-term consequences of mutation biases, helping to understand how the mutation process itself constrains sequence evolution.

## Methods

### Data

The 5-mer context mutation model was obtained from [16], where the mutability of each possible 5-mer sequence context was inferred from mutation events observed in reconstructed phylogenies of out-of-frame IgG lineages from [47]. For each mutation, the ancestral sequence was reconstructed to identify the 5-mer context present at the time of the mutation. The mutation rate at position *x* with context *w* was modeled as the product of a context-dependent mutability, *γ*_*w*_, and a position-dependent factor *β*_*x*_. The context- and position-dependent parameters were estimated jointly by maximizing the likelihood of the observed mutation events across reconstructed phylogenetic trees, using an iterative optimization procedure that separately updated the two sets of parameters. Because *γ*_*w*_ and *β*_*x*_ enter the model only through their product, their individual scales are not identifiable. To resolve this, the position-dependent factors were normalized such that ⟨*β*_*x*_⟩ = 1, fixing the relative scale of the two contributions. Then, *γ*_*w*_ > 1 indicates a 5-mer context with above-average mutability, whereas *γ*_*w*_ < 1 indicates a relatively “cold” context. For our analysis here, we used only the context-dependent factors, setting *m*_*w*_ = *γ*_*w*_ and treating all positions equivalently rather than incorporating *β*_*x*_.

To simulate somatic hypermutation using real BCR sequences, we randomly selected 2,000 sequences longer than 400 base pairs from the Observed Antibody Space database [23] (query: Species = human, Source = PBMC, BType = Naive-B-Cells, Chain = Heavy, Isotype = IGHM). To estimate changes in CDR3 amino acid frequencies before and after somatic hypermutation, we used the dataset from [28], comprising 7,228,250 naive and 8,521,052 memory BCR sequences from three individuals.

To simulate the somatic hypermutation effect on CDR1 and CDR2 across different V-gene families, we used germline V gene sequences from the ImMunoGeneTics (IMGT) database [29] (LIGM-DB query: Species = Homo sapiens (human), configuration_type = germline, gene_type = variable, functionality = functional, chain_type = IG-Heavy, selected labels: V-GENE, CDR1-IMGT, CDR2-IMGT). Because V-gene families contain different numbers of genes and their CDRs vary in length, we randomly sampled 100 9-mers for the Gillespie simulations from the list of all 9-mers derived from the CDR1/2 sequences within each family.

### Construction of the mutation rate matrix

For each sequence *i* of length *k*, we define *w*_*c*_(*i*) as a 5-nucleotide context centered at position *c*:

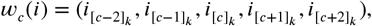

where

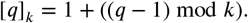

For example, for *k* = 9 and *c* = 1, *w*_*c*_(*i*) = (*i*_8_, *i*_9_, *i*_1_, *i*_2_, *i*_3_). This implements our “padding” procedure, equivalent to imposing periodic boundary conditions, by taking missing flanking nucleotides from the opposite end of the sequence.

For each nucleotide context *w*, the context model provides the mutability of its central nucleotide *m*_*w*_ and the probability of substituting it with any other nucleotide 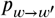, given that mutation occurs. For two different sequences *i* and *j*, we define the mutation rate from *i* to *j* as:

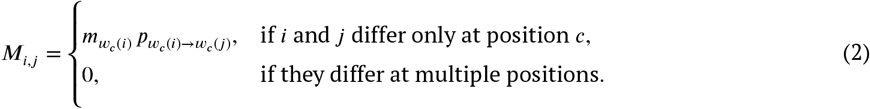

The total mutability of sequence *i* is: 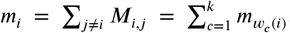. The diagonal entries are then defined as *M*_*i,i*_ = −*m*_*i*_, so that each row of *M* sums to zero.

For the main analysis, we used *k* = 9, resulting in a mutation rate matrix with 4^9^ = 262, 144 sequences. As a result, the mutation rate matrix was sparse, with non-zero off-diagonal entries only to 3*k* = 27 other sequences differing by a single-nucleotide substitution.

### Gillespie simulations

Gillespie simulations on k-mers were performed by treating each k-mer as a state in a mutation Markov process. The waiting time to the next mutation was sampled from an exponential distribution with the rate equal to the outgoing mutation rate, and the next k-mer (differing by a single substitution) was selected with probability proportional to the corresponding rate in *M*. This process was repeated until a specified stopping time across multiple replicate trajectories. We then pooled the resulting trajectories and recorded the number of times each k-mer was visited.

Gillespie simulations on BCR sequences were performed by dividing each sequence into overlapping 5-mers using a sliding window. At each simulation step, a 5-mer was selected for mutation with probability proportional to its mutability from [16] (see description in *Methods: Data*), multiplied by the number of times it appears in the current sequence. The waiting time to the next event was sampled from an exponential distribution using the total sequence-wide mutation rate, equal to the sum of 5-mer mutabilities, weighted by their number of occurrences in the sequence. The new nucleotide was then chosen according to the 5-mer model of somatic hypermutation [16], and the affected overlapping 5-mers were updated. We ran this simulation for a time substantially exceeding the characteristic timescale of relaxation towards the stationary distribution, after which we measured 5-mer frequencies in the resulting sequence.

### Codon table simulations

We performed codon-table simulations preserving the number of codons assigned to each amino acid, as well as the internal structure of each synonymous codon set, including the Hamming-distance relationships among codons. To achieve that, we generated alternative genetic codes by independently permuting the nucleotide alphabet at each of the three codon positions and applying the same position-specific mappings to all 64 codons. This procedure preserves the codon blocks assigned to each amino acid while randomizing the nucleotide identities within that block. We enumerated all 24^3^ = 13, 824 possible combinations of permutations. For each resulting genetic code, we calculated the stationary frequency of each amino acid by summing the stationary frequencies of its assigned codons (obtained by aggregating the stationary frequencies of all 9-mers sharing the corresponding central codon).

## Author contributions

Conceptualization: VAK, DMM; Methodology: VAK, DMM; Formal analysis: VAK, DMM; Writing - original draft: VAK, DMM; Writing - review & editing: VAK, HVM, DMM; Visualization: VAK; Supervision: HVM, DMM.

## Code availability

Custom analysis code was written in Python (version ≥ 3.10.9). All analysis code is freely available on GitHub: https://github.com/meyer-lab-cshl/SHM-space_paper.

## Supplementary Figures

**Extended Data Figure 1.**
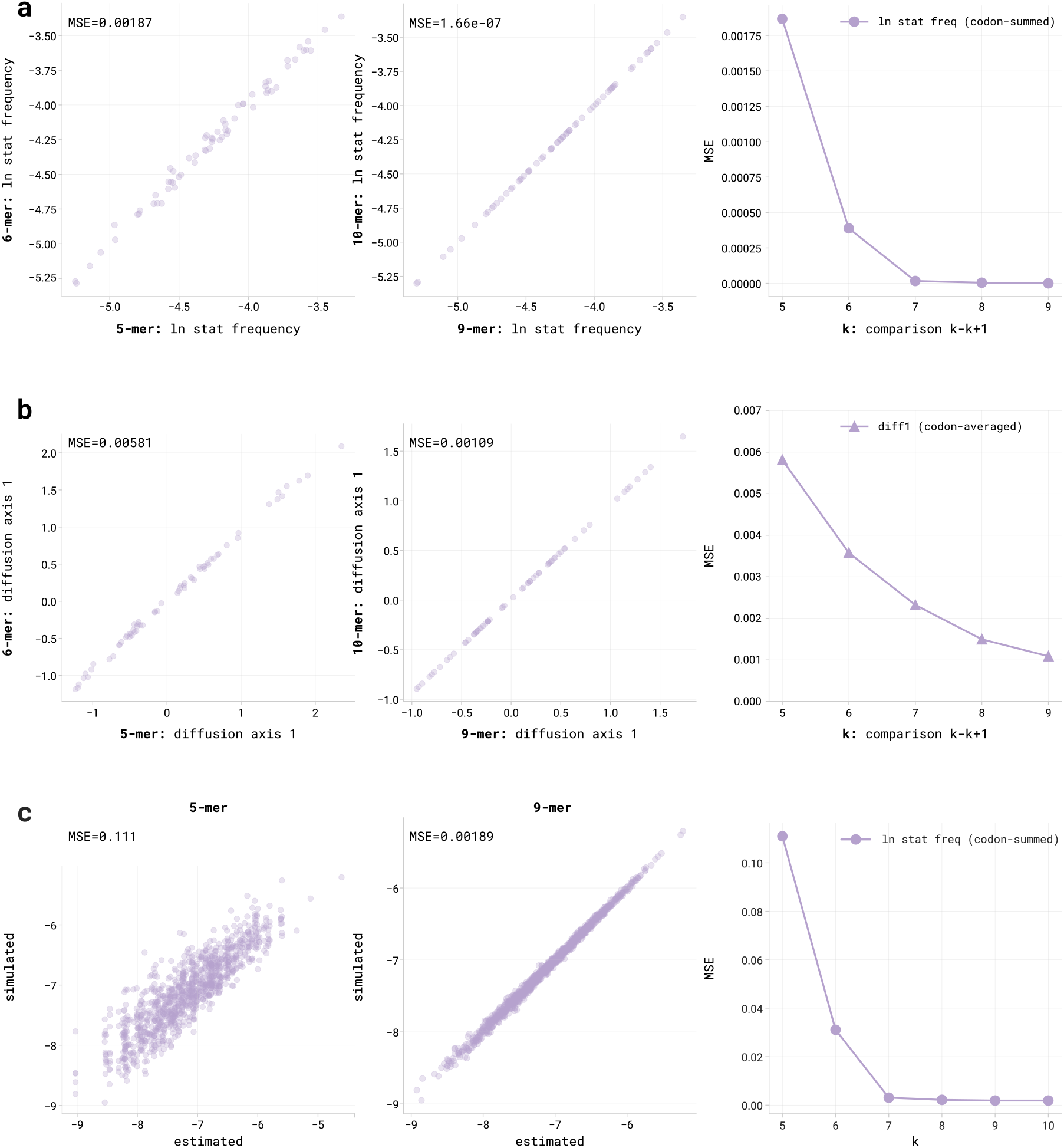
Validation of the padding procedure and selection of the k-mer length. **(a)** Comparison of codon-level log stationary frequencies calculated from mutation matrices of consecutive *k*-mer lengths. Scatter plots show the correspondence between 5-mers and 6-mers (left) and between 9-mers and 10-mers (middle). Right: mean squared error (MSE) between consecutive *k*-mer lengths for *k* = 5 … 9. **(b)** The same comparisons as in (a), but for diffusion axis 1. **(c)** Comparison of log stationary frequencies estimated from the padded k-mer mutation matrices and those obtained from mutation simulations on full-length BCR sequences. Scatter plots show results for 5-mers (left) and 9-mers (middle). Right: MSE between estimated and simulated log stationary frequencies for *k* = 5 … 10.

**Extended Data Figure 2.**
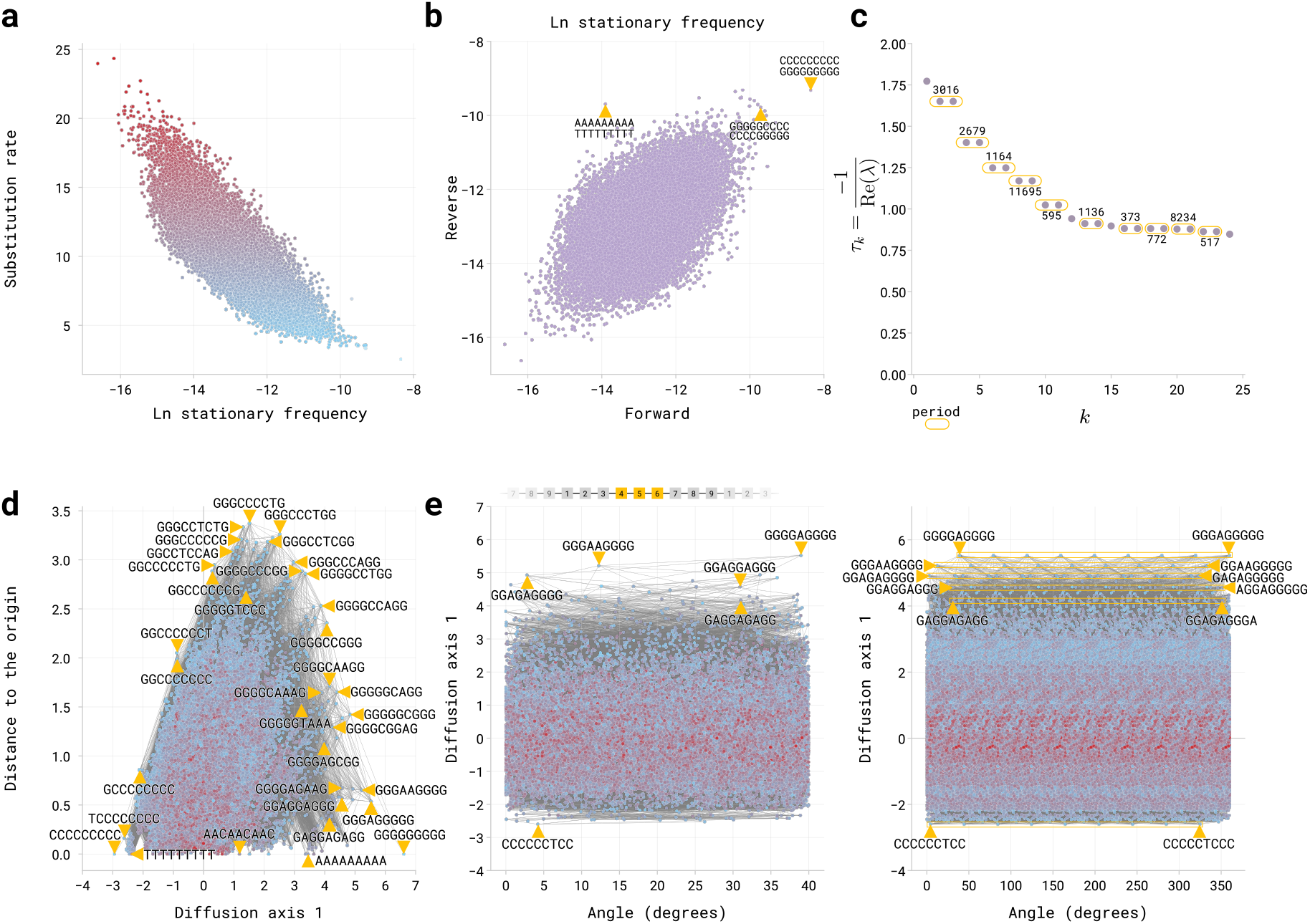
Additional properties of the 9-mer mutational landscape. **(a)** Relationship between the substitution rate and log stationary frequencies of 9-mers (colored by substitution rate: red = high, blue = low). **(b)** Comparison of log stationary frequencies for 9-mers that are reverse complements of each other. **(c)** Characteristic decay times 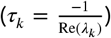 of deviations from stationarity plotted against eigenspace index (*k*). Numbers indicate the corresponding oscillation periods for complex eigenvalues. **(d)** Distance from the origin in Figure 1c plotted against diffusion axis 1. **(e)** Angular coordinate from Figure 1c plotted against the diffusion axis 1, showing one segment of angular range (left, 40^°^) or the full range (right, 360^°^). Lines connect sequences differing by a single-nucleotide substitution.

**Extended Data Figure 3.**
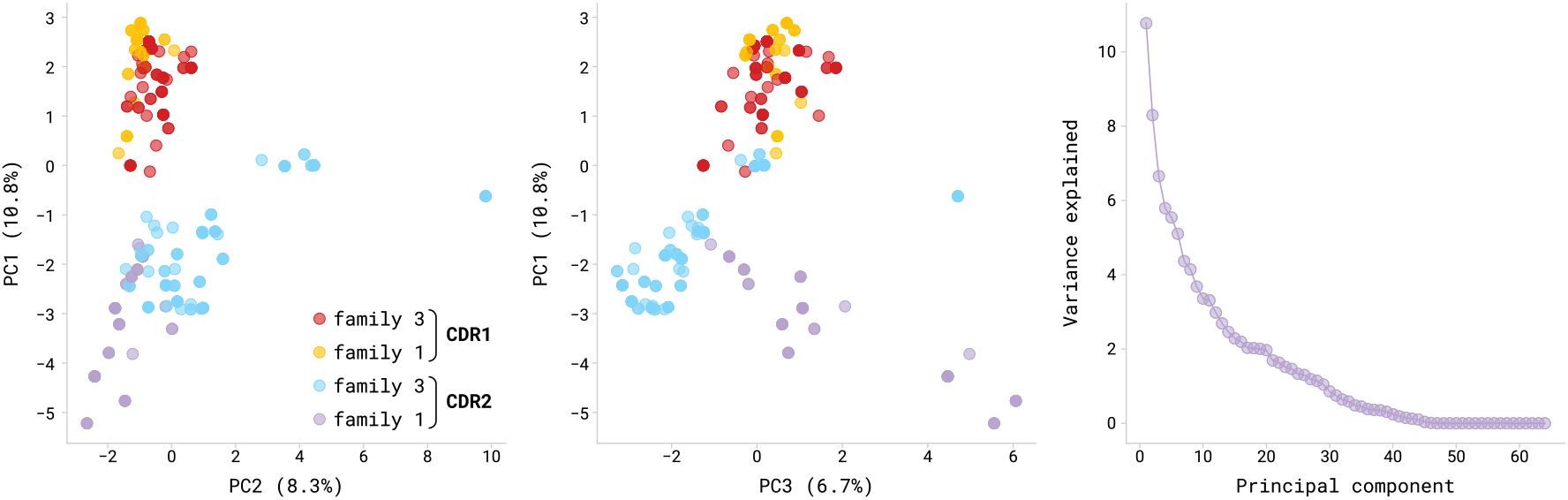
Projections of amino acid frequencies of CDR1 and CDR2 sequences of V-genes from the IGHV1 and IGHV3 families onto the first and second principal components (left) and the first and third principal components (middle). Each point represents a CDR sequence, colored by a gene family and a CDR type. Right: variance explained by each principal component.

